# Shifts in colour morph frequencies along an urbanisation gradient in the ground beetle *Pterostichus madidus*

**DOI:** 10.1101/2023.03.31.535151

**Authors:** Maxime Dahirel, Hélène Audusseau, Solène Croci

## Abstract

Cities impose strong ecological constraints on many species. It is often difficult to know whether phenotypic responses to urbanisation are due to (adaptive) evolutionary responses, as opposed to e.g. plastic responses. A solution is to focus on traits for which variation is only or mostly genetic in origin, since changes are then likely to reflect evolutionary processes. For this purpose, we studied the leg-colour polymorphism of a common carabid beetle, *Pterostichus madidus*, along an urbanisation gradient. We observed that black-legged individuals were more frequent than red-legged individuals in urban areas. Whether these changes result from natural selection or non-selective evolutionary forces is uncertain. However, if this phenotypic change is adaptive in nature, higher urban temperatures are likely to be the driver. Specifically, our results are consistent with previous data showing that black-legged individuals have a behavioural advantage in warmer (micro)climates, and contradict the thermal melanism hypothesis that predicts they would experience stronger negative effects of higher temperatures in cities.

## Introduction

Although cities occupy a small part of the planet, their continued expansion (Seto et al. 2012) raises key questions about their impact on biodiversity. While community-level responses to urbanisation have been the focus of much interest (e.g. Merckx et al. 2018, Piano et al. 2020), and within-species trait variation along urbanisation gradients and mosaics has been repeatedly reported (Alberti et al. 2017), examples of urban phenotypic changes unambiguously attributable to evolution are rare (Lambert et al. 2021). The data requirements for clear demonstrations of urban evolution can be challenging, as observed trait variability may not necessarily result from evolutionary changes (Lambert et al. 2021). One solution is to study traits for which variation is overwhelmingly genetic in origin with little to no phenotypic plasticity. In that case, phenotypic changes are by definition most likely attributable to evolutionary processes, whether adaptive or not. For instance in the white clover (*Trifolium repens*), the ability to produce hydrogen cyanide is controlled by two loci and can easily be evaluated on wild plants, which facilitated the study of its urban evolution at a global scale (Santangelo et al. 2022).

The black clock beetle *Pterostichus madidus* (Fabricius, 1775)(fam. Carabidae) may be one of these species particularly well-suited to urban evolution studies. It is one of the most abundant carabids in both urban and non-urban woodlands in (north)western Europe (Sadler et al. 2006; Croci et al. 2008). Individuals exhibit colour variation, having either red or black legs (**Fig. 1**), whose inheritance seems to obey simple Mendelian rules (Pudney 2002). Morph frequencies vary substantially among sites, including over short distances (Terrell-Nield 1990; Pudney 2002). Two mechanisms could explain these frequency variations, leading to opposite predictions about how leg colour responds to urbanisation, specifically to the higher temperatures observed in cities (Foissard et al. 2019). First, red-legged *P. madidus* seem to have their activity peak at lower temperatures than black-legged individuals, and may be active under a narrower humidity range (Fairhurst 1969). This may explain why, according to British studies on spatial patterns in *P. madidus* morph frequencies, red-legged morphs are more frequent in colder and oceanic landscapes than hotter and drier ones, and in the cooler inner part of woodlands than outside (Terrell-Nield 1990; Pudney 2002). Based on this, we would expect red-legged beetles to be rarer in warmer urban contexts. Second, under the thermal melanism hypothesis, darker forms should typically suffer more from warm (micro)climates as they heat faster than lighter forms (Clusella Trullas et al. 2007). We would then expect black-legged beetles to be disadvantaged and rarer in cities. To discriminate between these hypotheses, we revisited data from a previously published community ecology study (Croci et al. 2008), in which morph information happened to be collected for *P. madidus*.

**Figure 1.**
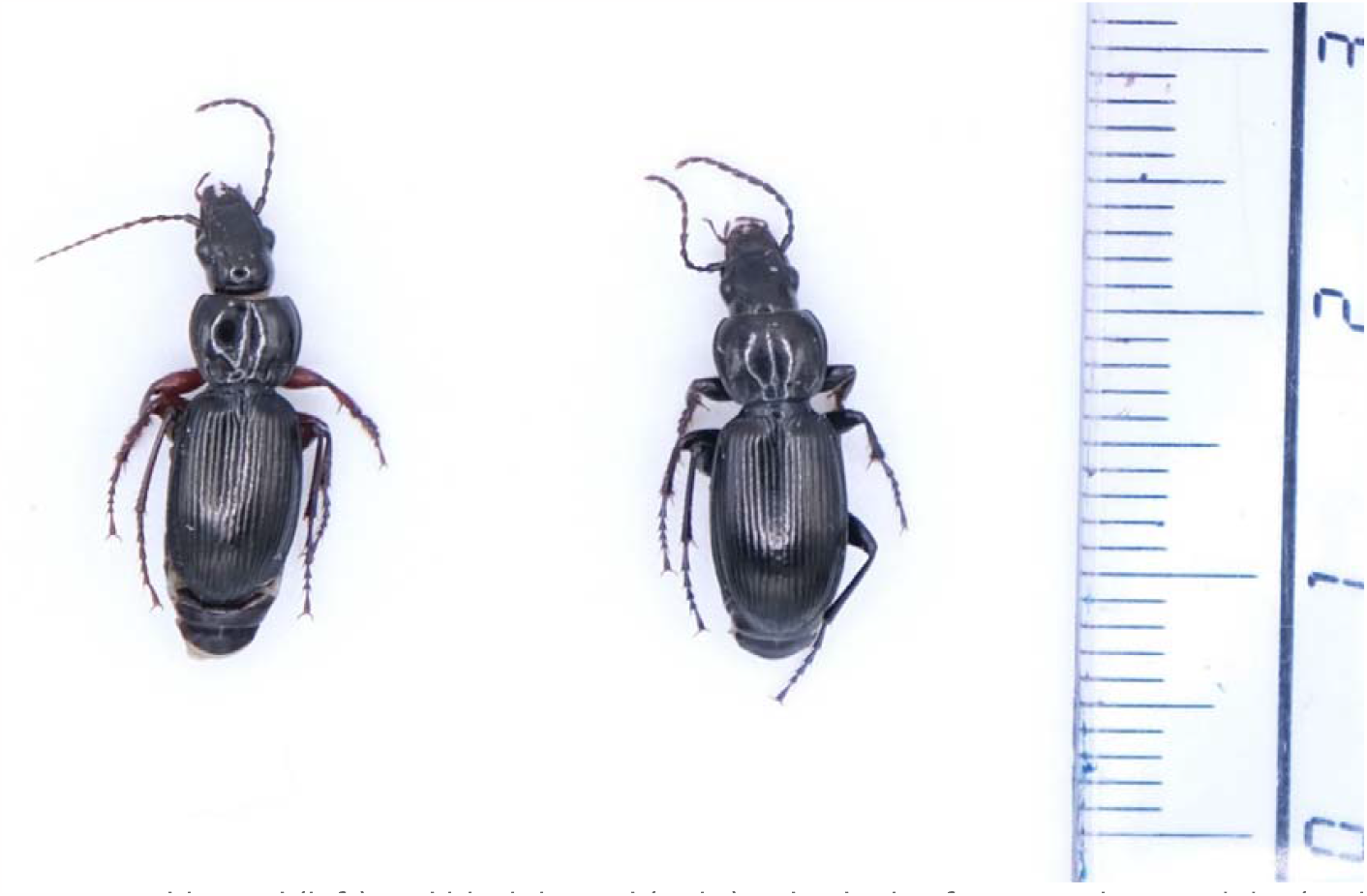
Red-legged (left) and black-legged (right) individuals of *Pterostichus madidus* (scale in cm). Cropped photograph, see **Data availability** for original image.

## Methods

### Site selection and sampling

The data reanalysed here are from Croci et al. (2008), with the addition of previously unused data from two sites sampled over the same period. Beetles were collected in 2004 and 2005 over 14 small woodlands in and around Rennes (western France, **Fig. 2**). In each site, 12 pitfall traps were installed, filled with 30% ethanol and covered to protect them from rain and litter. Traps were emptied roughly every two weeks from late March to early November 2004 and again from mid-April to early August 2005, for a total of 24 sampling sessions. A total of 9934 *P. madidus* individuals were collected and their leg colour recorded (**Data availability**). In the present study, beetles from traps from the same woodland × sampling session combination are pooled together.

**Figure 2.**
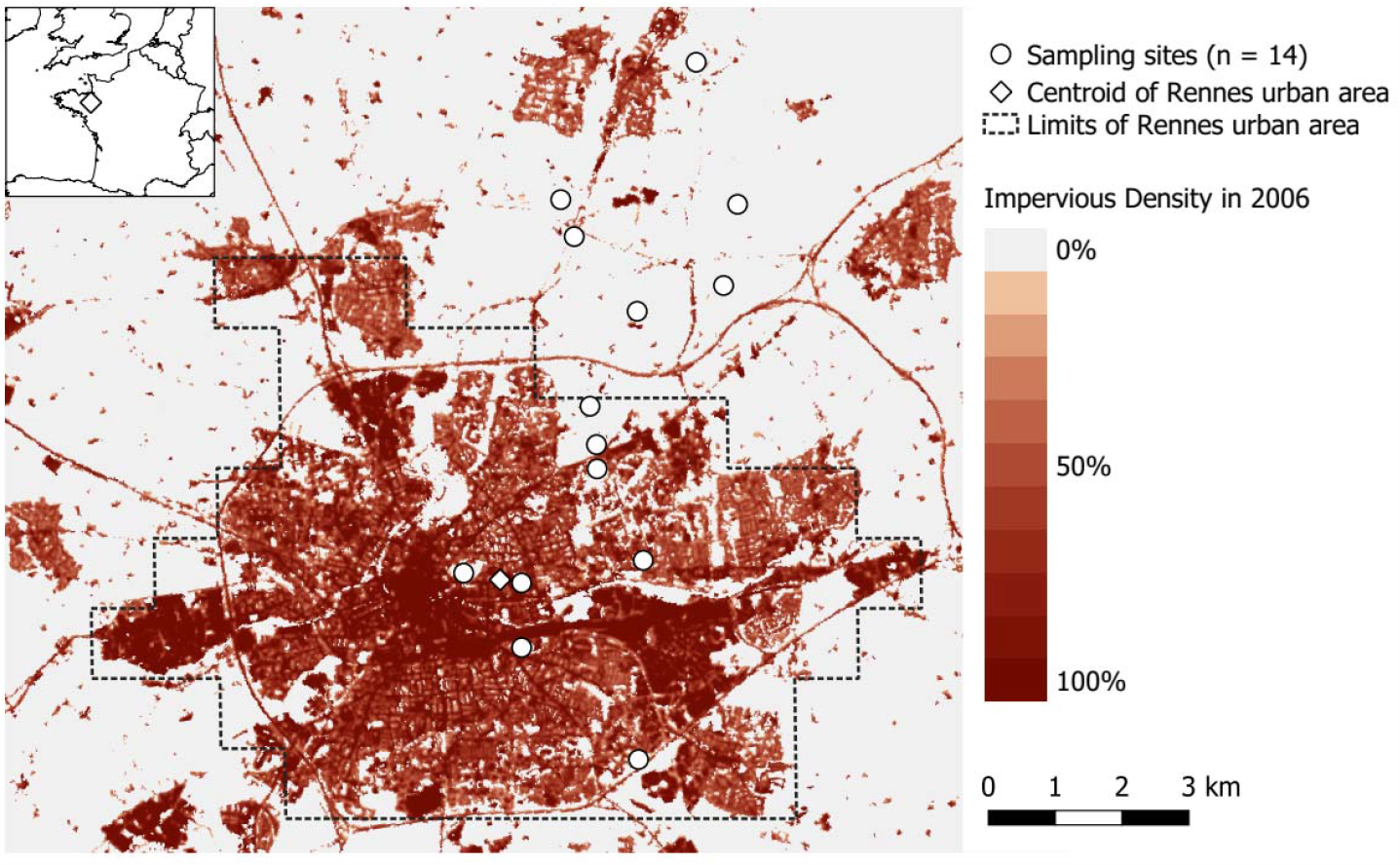
Map of Rennes (western France) showing the location of the studied woodlands, the boundary of the Rennes urban area and its centroid.

### Urbanisation metrics

We assessed urbanisation in two ways using remote sensing-derived layers. First, we used the distance between the centroids of the studied woodlands and the centroid of the Rennes urban area, taken from the GHS Urban Centre Database (**Fig. 2**, v1.2, Florczyk et al. 2019). Second, we used the average imperviousness around each woodland, based on the Imperviousness Density High Resolution Layer (**Fig. 2**, 2006 version, European Environmental Agency 2018). This layer was chosen over other data sources because of its fine resolution (20 m) and temporal proximity to the sampling years. We explored the effects of spatial scale by estimating mean Imperviousness Density within buffers of 100, 300, 600, 900, 1200, 1500 and 1800 m radius around the centroid of each woodland (the scales range was based on Croci et al. 2008, Merckx et al. 2018, and Piano et al. 2020).

### Statistical analyses

We analysed data in a Bayesian framework using R (version 4.2.2, R Core Team 2022) and the *brms* package (Bürkner 2017).

We built beta-binomial generalised linear mixed models with the proportion of black-legged beetles per woodland and sampling session as the response (with a logit link, see **Supplementary Material S1** for a detailed model description). These models included a fixed effect of urbanisation, a random intercept effect of site identity to account for repeated measurements at each site, and random effects of sampling session (random intercept and random urbanisation slope) to account for consistent temporal variation across all sites. We built one model per urbanisation metric (distance to city centroid, and imperviousness at various scales). Models were compared both based on their overall predictive performance using K-fold cross-validation, and based on the proportion of among-site variance explained by urbanisation (see **Supplementary Material S2**).

Urbanisation metrics were scaled to mean 0 and unit 1SD at the site level to facilitate model estimation and prior setting. We chose weakly informative priors (**Supplementary Material S1**) and ran four chains per model for 2000 iterations each, with the first half of each chain used as warmup. All parameters had satisfactory effective sample sizes and convergence (Vehtari et al. 2021). Posteriors are summarized as means [95% Highest Posterior Density Intervals].

## Results

All models performed very similarly and led to the same qualitative interpretations (**Supplementary Material S2** and **S3**). We only present below results for the model using Imperviousness Density in a 100 m radius (the one with the highest proportion of among-site variance explained by urbanisation) for simplicity; the other models are detailed in **Supplementary Material S3**.

The proportion of black-legged beetles increased with urbanisation (**Fig. 3**, fixed effect slope of imperviousness: = 0.50 [0.30; 0.69]). This slope shows little variation across sampling sessions (the SD of the urbanisation random effect slopes is 10.85% [0.03%; 27.30%] of the absolute value of the fixed effect slope; **Supplementary Material S4**).

**Figure 3.**
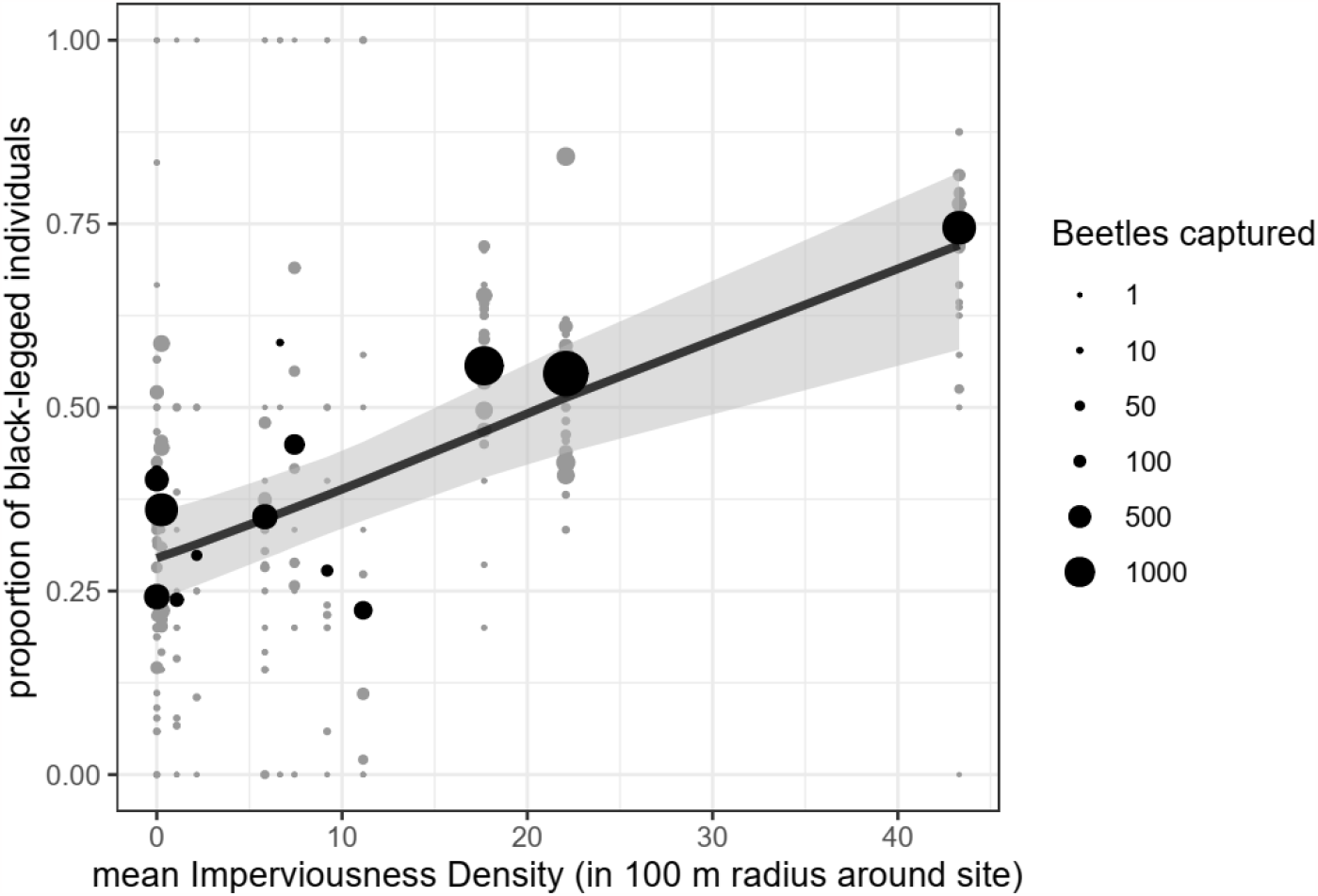
Proportion of *Pterostichus madidus* beetles with black legs as a function of urbanisation (Imperviousness Density in 100 m radius). Grey dots: observed proportions for each site × sampling session combination, black dots: site-level weighted averages across all sampling sessions, black line and grey bands: mean predicted relationship and 95% credible band.

## Discussion

By revisiting an 18 years old dataset, we showed that the frequency of black-legged *Pterostichus madidus* individuals increased along an urbanisation gradient, rejecting – at least locally – the hypothesis that thermal melanism (predicting fewer dark forms in warm microclimates) shapes urban morph frequencies in this species.

Behavioural differences in morph responses to climatic conditions (Fairhurst 1969) could lead to variation in trappability along the urbanisation gradient, and thus explain the observed shift in morph data, *irrespective of the actual morph frequencies* (Álvarez-Quintero et al. 2021). However, given the lack of seasonal variation in morph frequency and its response to urbanisation, observed morph shifts more likely reflect true morph shifts.

Under the basis that leg colour is genetically determined (Pudney 2002), the shift in morph frequencies observed along the urbanisation gradient is most likely evolutionary. Whether it results from natural selection is a more difficult question. Non-selective forces such as genetic drift can lead to such evolutionary clines (Santangelo et al. 2018). However, while we cannot fully exclude them using our data, they are unlikely to play a major role here. For instance regarding drift, genetic data from *P. madidus* in other cities (Desender et al. 2005) show no consistent loss of neutral genetic diversity in urban sites, and we found no strong evidence for shifts in population sizes along the gradient (**Supplementary Material S5**).

Temperature is likely a major selective force underlying the variation in morph frequencies. Urban areas are often characterized by higher temperatures, as is the case in our studied city (Foissard et al. 2019). Consistently, other studies also found a correlation between warmer climate and higher frequencies of black-legged individuals (Terrell-Nield 1990; Pudney 2002). Still, the observed pattern could be caused by another of the many environmental factors altered by urbanisation (Parris 2016). Multi-city comparisons may help here, as cities vary in the intensity of various environmental drivers (see e.g. Santangelo et al. 2022). To our knowledge, information on urban *P. madidus* morphs is available for one other city (Terrell-Nield 1990; Pudney 2002), but the coarser scale of urbanisation data in these older studies prevents direct comparison. In any case, this work highlights the potential value for urban evolution of ecological studies recording morphs of conspicuously polymorphic species. Further studies may also help identify the underlying mechanisms behind this evolutionary response in more detail. The link between colour and urban environmental change may even not be direct as melanin and melanin synthesis pathways have wide-ranging effects including on immune function, UV tolerance, and mechanical resistance (San-Jose and Roulin 2018). Mechanistic and physiological studies of their response to urbanisation would certainly be enlightening.

## Acknowledgments

We are grateful to the many people involved in the original study, including Patricia Le Quilliec, Anita Le Brecht-Georges, Geoffrey Desjardins, Rose-Anne Fracy, Philippe Clergeau and Alain Butet. We also thank the city of Rennes and Rennes Métropole for funding and supporting the original data collection through the Ecorurb programme.

## Data availability

R code and data needed to reproduce the analyses in this manuscript are accessible from Github (https://github.com/mdahirel/pterostichus-morphology-2004) and archived in Zenodo (https://doi.org/10.5281/zenodo.7737152).

## Author contributions

Initial idea: MD, HA and SC; contributed and curated initial data: SC; analysed data: MD; wrote the initial draft: MD. All authors revised and edited the manuscript.

## References

Alberti M, Correa C, Marzluff JM, et al (2017) Global urban signatures of phenotypic change in animal and plant populations. PNAS 201606034. https://doi.org/10.1073/pnas.1606034114

Álvarez-Quintero N, Chiara V, Kim S-Y (2021) Trap versus net: Behavioural sampling bias caused by capture method in three-spined sticklebacks. Behav Processes 193:104504. https://doi.org/10.1016/j.beproc.2021.104504

Bürkner P-C (2017) brms: an R package for Bayesian multilevel models using Stan. J Stat Softw 80:1–28. https://doi.org/10.18637/jss.v080.i01

Clusella Trullas S, van Wyk JH, Spotila JR (2007) Thermal melanism in ectotherms. J Therm Biol 32:235–245. https://doi.org/10.1016/j.jtherbio.2007.01.013

Croci S, Butet A, Georges A, et al (2008) Small urban woodlands as biodiversity conservation hot-spot: a multi-taxon approach. Landscape Ecol 23:1171–1186. https://doi.org/10.1007/s10980-008-9257-0

Desender K, Small E, Gaublomme E, Verdyck P (2005) Rural-urban gradients and the population genetic structure of woodland ground beetles. Conserv Genet 6:51–62. https://doi.org/10.1007/s10592-004-7748-3

European Environmental Agency (2018) High Resolution Layer: Imperviousness Degree (IMD) 2006

Fairhurst JM (1969) Aspects of activity and density in some Carabidae. PhD thesis, University of Manchester

Florczyk AJ, Melchiorri M, Corbane C, et al (2019) Description of the GHS Urban Centre Database 2015. Publications Office of the European Union, Luxembourg. https://doi.org/10.2760/037310

Foissard X, Dubreuil V, Quénol H (2019) Defining scales of the land use effect to map the urban heat island in a mid-size European city: Rennes (France). Urban Climate 29:100490. https://doi.org/10.1016/j.uclim.2019.100490

Lambert MR, Brans KI, Des Roches S, et al (2021) Adaptive evolution in cities: progress and misconceptions. Trends Ecol Evol 36:239–257. https://doi.org/10.1016/j.tree.2020.11.002

Merckx T, Souffreau C, Kaiser A, et al (2018) Body-size shifts in aquatic and terrestrial urban communities. Nature 558:113–116. https://doi.org/10.1038/s41586-018-0140-0

Parris KM (2016) Ecology of urban environments. Wiley-Blackwell, Chichester, UK

Piano E, Souffreau C, Merckx T, et al (2020) Urbanization drives cross-taxon declines in abundance and diversity at multiple spatial scales. Glob Chang Biol 26:1196–1211. https://doi.org/10.1111/gcb.14934

Pudney K (2002) Investigation of leg colour polymorphism in Pterostichus madidus (F.) in relation to climatic factors. PhD thesis, Nottingham Trent University

R Core Team (2022) R: a language and environment for statistical computing

Sadler JP, Small EC, Fiszpan H, et al (2006) Investigating environmental variation and landscape characteristics of an urban–rural gradient using woodland carabid assemblages. J Biogeogr 33:1126–1138. https://doi.org/10.1111/j.1365-2699.2006.01476.x

San-Jose LM, Roulin A (2018) Toward understanding the repeated occurrence of associations between melanin-based coloration and multiple phenotypes. Am Nat 192:111–130. https://doi.org/10.1086/698010

Santangelo JS, Johnson MTJ, Ness RW (2018) Modern spandrels: the roles of genetic drift, gene flow and natural selection in the evolution of parallel clines. Proc Royal Soc B 285:20180230. https://doi.org/10.1098/rspb.2018.0230

Santangelo JS, Ness RW, Cohan B, et al (2022) Global urban environmental change drives adaptation in white clover. Science 375:1275–1281. https://doi.org/10.1126/science.abk0989

Seto KC, Güneralp B, Hutyra LR (2012) Global forecasts of urban expansion to 2030 and direct impacts on biodiversity and carbon pools. PNAS 109:16083–16088. https://doi.org/10.1073/pnas.1211658109

Terrell-Nield CE (1990) Distribution of leg-colour morphs of Pterostichus madidus (F.) in relation to climate. In: Stork NE (ed) The role of ground beetles in ecological and environmental studies. Intercept, Andover, pp 39–51

Vehtari A, Gelman A, Simpson D, et al (2021) Rank-normalization, folding, and localization: an improved R for assessing convergence of MCMC (with discussion). Bayesian Anal 16:667–718. https://doi.org/10.1214/20-BA1221

