## Supplementary Material for "Shifts in colour morph frequencies along an urbanisation gradient in the ground beetle *Pterostichus madidus*"

#### S1 - Model description

For the present study, our response variable is $B_{i,j}/N_{i,j}$, the proportion of black-legged *P. madidus* beetles captured in a given woodland $i$ during a given sampling session $j$, with $B_{i,j}$ the number of black-legged beetles and $N_{i,j}$ the number of individuals of both morphs. Note that we naturally only include woodland × session combinations with $N_{i,j} > 0$. We ran the models below for each of the 8 possible urbanisation metrics described in the main text.

We initially built binomial models as follows:

$$B_{i,j} \sim \text{Binomial}(N_{i,j}, p_{i,j}),$$
$$\text{logit}(p_{i,j}) = \beta_0 + (\beta_1 + \eta_j) \times x_i + \alpha_i + \gamma_j,$$

with $x_i$ the (centered and scaled) urbanisation metric at site $i$, $\beta_0$ and $\beta_1$ the fixed effects (intercept and urbanisation slope respectively), $\alpha_i$ the site-specific random-effect intercept, and $\gamma_j$ and $\eta_j$ the session-specific random intercept and slope. Random effects are distributed as follows:

$$\alpha_i \sim \text{Normal}(0, \sigma_\alpha),$$
$$\begin{bmatrix}\gamma_j \\ \eta_j\end{bmatrix} \sim \text{MVNormal}\left(\begin{bmatrix}0 \\ 0\end{bmatrix}, \mathbf{\Omega}\right),$$

where $\mathbf{\Omega}$ is the covariance matrix for the session-specific random effects, which can be decomposed into its constituent standard deviations and correlation matrix $\boldsymbol{R}$:

$$\mathbf{\Omega} = \begin{bmatrix}\sigma_\gamma & 0 \\ 0 & \sigma_\eta\end{bmatrix} \boldsymbol{R} \begin{bmatrix}\sigma_\gamma & 0 \\ 0 & \sigma_\eta\end{bmatrix}.$$

We used weakly informative priors mostly inspired by McElreath (2020). We used $\text{Normal}(0, 1.5)$ priors for the fixed effect intercept $\beta_0$, which corresponds to the logit of a proportion, and $\text{Normal}(0, 1)$ for the fixed effect slope $\beta_1$. We used $\text{Half} - \text{Normal}(0, 1)$ priors for all standard deviations $\sigma$, and a $\text{LKJ}(2)$ prior for the correlation matrix $\boldsymbol{R}$.

Evaluations of these models revealed slight but consistent evidence of overdispersion (see archived data analysis code: https://github.com/mdahirel/pterostichus-morphology-2004 or https://doi.org/10.5281/zenodo.7737152). We therefore fitted beta-binomial models to account for that overdispersion (Harrison 2015):

$$B_{i,j} \sim \text{BetaBinomial}(N_{i,j}, p_{i,j}, \phi),$$

which is equivalent to

$$B_{i,j} \sim \text{Binomial}(N_{i,j}, \pi_{i,j}),$$
$$\pi_{i,j} \sim \text{Beta}(a_{i,j}, b_{i,j})$$
$$a_{i,j} = p_{i,j} \times \phi$$
$$b_{i,j} = (1 - p_{i,j}) \times \phi,$$

where $\phi$ is a precision parameter (higher values = lower overdispersion) in the `brms` (Bürkner 2017) implementation. We set priors on the inverse of $\phi$ (where higher values = higher overdispersion): $1/\phi \sim \text{Half} - \text{Normal}(0, 1)$ (based on similar suggestions for negative binomial models; seee e.g. https://github.com/stan-dev/stan/wiki/Prior-Choice-Recommendations#story-when-the-generic-prior-fails-the-case-of-the-negative-binomial). The model formulation and priors for $p_{i,j}$ are then the same as in the binomial case.

### S2 - Model performance comparisons

We used two metrics to compare our models (both binomial and beta-binomial, although we only present beta-binomial models here for simplicity). We first used $K$-fold cross-validation (with $K = 10$) to evaluate these models based on their overall pointwise predictive accuracy (Vehtari et al. 2017). We then also compared them based on the proportion of (logit scale) among-site variance explained by fixed effects $\frac{\sigma^2_\beta}{\sigma^2_\beta + \sigma^2_\alpha}$, where the variance explained by fixed effects $\sigma^2_\beta$ is estimated as in Nakagawa and Schielzeth (2013). This is because since all models include a site random effect, comparing them on their overall performance may not reflect the way urbanisation specifically explains average among-site differences, as the site random effect will "absorb" any among-site variation not explained by the urbanisation metric.

All eight models had functionally identical overall predictive performance, based on cross-validation results (**Table S2-1**). The proportion of among-site variance explained by the effect of urbanisation was highest at the 100 m scale and decreased as buffer size increased, with the lowest value for the fixed effect of distance to the urban centroid (**Fig. S2-1**). However, there was actually limited support for differences between models, with posteriors for the proportion of site-level variance explained all largely overlapping (**Fig. S2-1**).

**Table S2-1.** Expected log pointwise predictive density (elpd) for each of the 8 beta-binomial models, expressed as differences ± SE from the elpd of the "best" model in the set.

| Urbanisation metric | elpd deviation from "best" model ± SE |
|---|---|
| IMD (600m) | 0.00 ± 0.00 |
| IMD (100m) | -0.30 ± 1.52 |
| IMD (900m) | -0.32 ± 0.49 |
| IMD (1200m) | -0.37 ± 0.44 |
| IMD (1500m) | -0.44 ± 0.52 |
| IMD (300m) | -0.52 ± 0.47 |
| IMD (1800m) | -0.52 ± 0.55 |
| distance to urban centroid | -1.67 ± 1.29 |

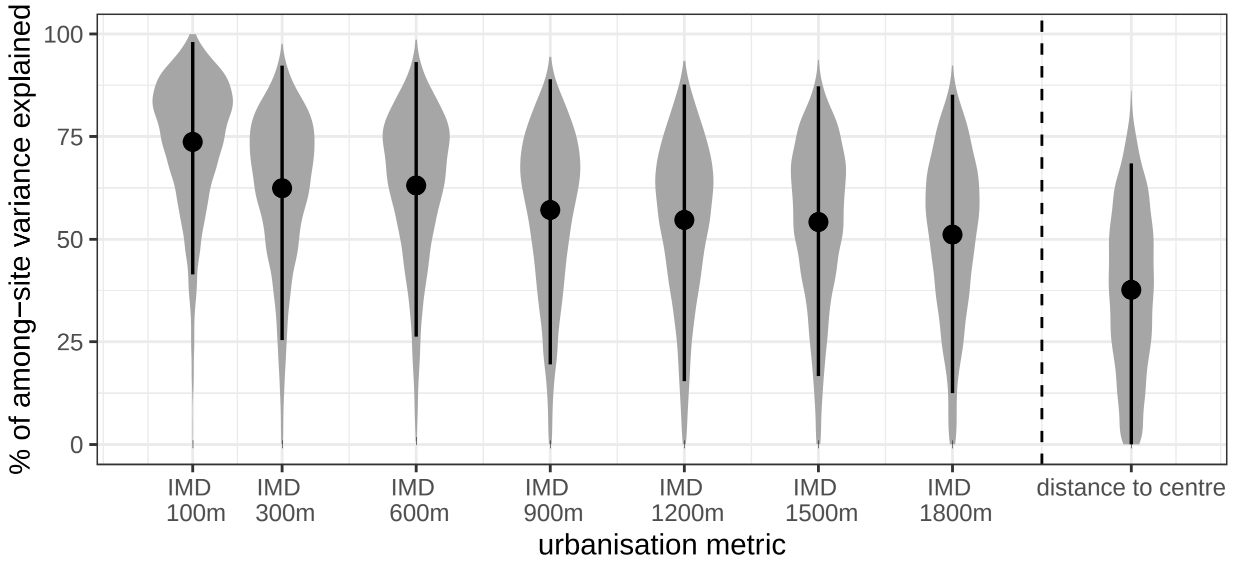


**Figure S2-1.** Posterior proportion of (logit-scale) site-level variance in morph frequencies explained by urbanisation, depending on the urbanisation metric used in the model. All models are the beta-binomial implementations; black dots are posterior means, segments 95% Highest Posterior Density Intervals.

### S3 - Comparing the effect of urbanisation across models

The standardised effect of urbanisation $\beta_1$ is quite consistent whether distance to city centroid or local Imperviousness Density is used as urbanisation metric, and regardless of the spatial scale at which Imperviousness Density is estimated (**Fig. S3-1**).

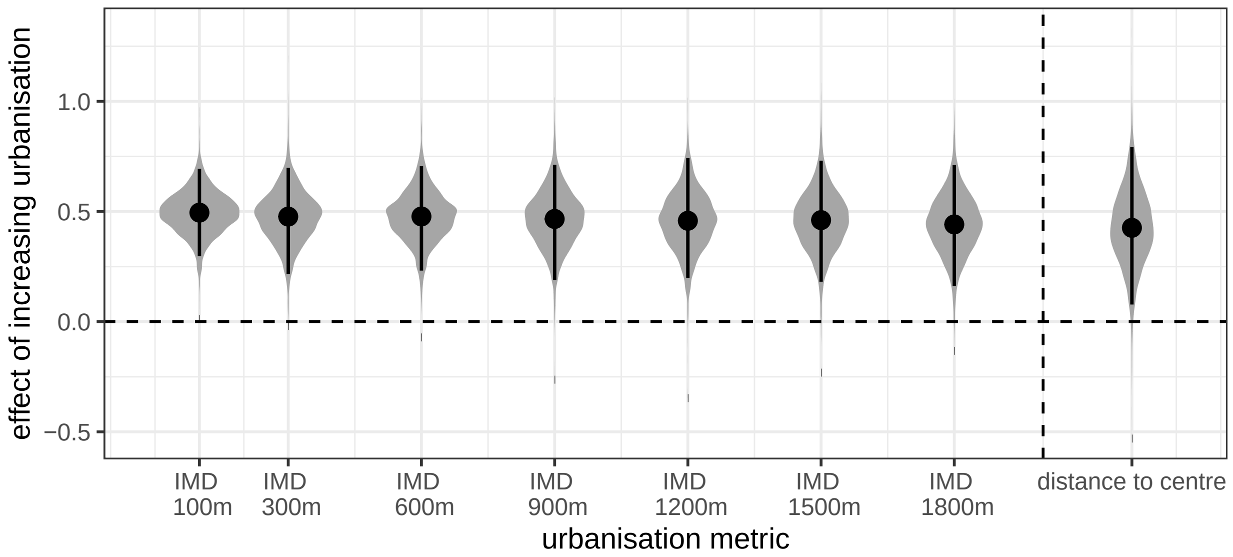


**Figure S3-1.** Posterior estimates for the standardised fixed effect of urbanisation $\beta_1$ on the proportion of black-legged *P. madidus*, depending on the urbanisation metric used in the model. Note that the sign of the posterior values for distance to city centroid has been reversed to make the comparison with the other posteriors easier (as distances to city centroid decrease when urbanisation increases). Black dots are posterior means, segments 95% Highest Posterior Density Intervals.

#### S4 - Seasonal variation in urbanisation effect

As discussed in the main text, between-session variation in the effect of urbanisation is small. This is a strong hint that our results reflect true morph differences and not simply weather-induced seasonal variation in relative morph activity (and trappability).

This seasonal stability can be seen by plotting the random effects to see how the urbanisation effect deviates from the overall mean effect session by session (random slopes, **Fig. S4-1, top**). One can also do the same for the session-specific random intercepts (**Fig. S4-1, bottom**), or by visualising the consequences of both in terms of session-specific predicted values (**Fig. S4-2**). In all cases, no obvious seasonal pattern can be seen.

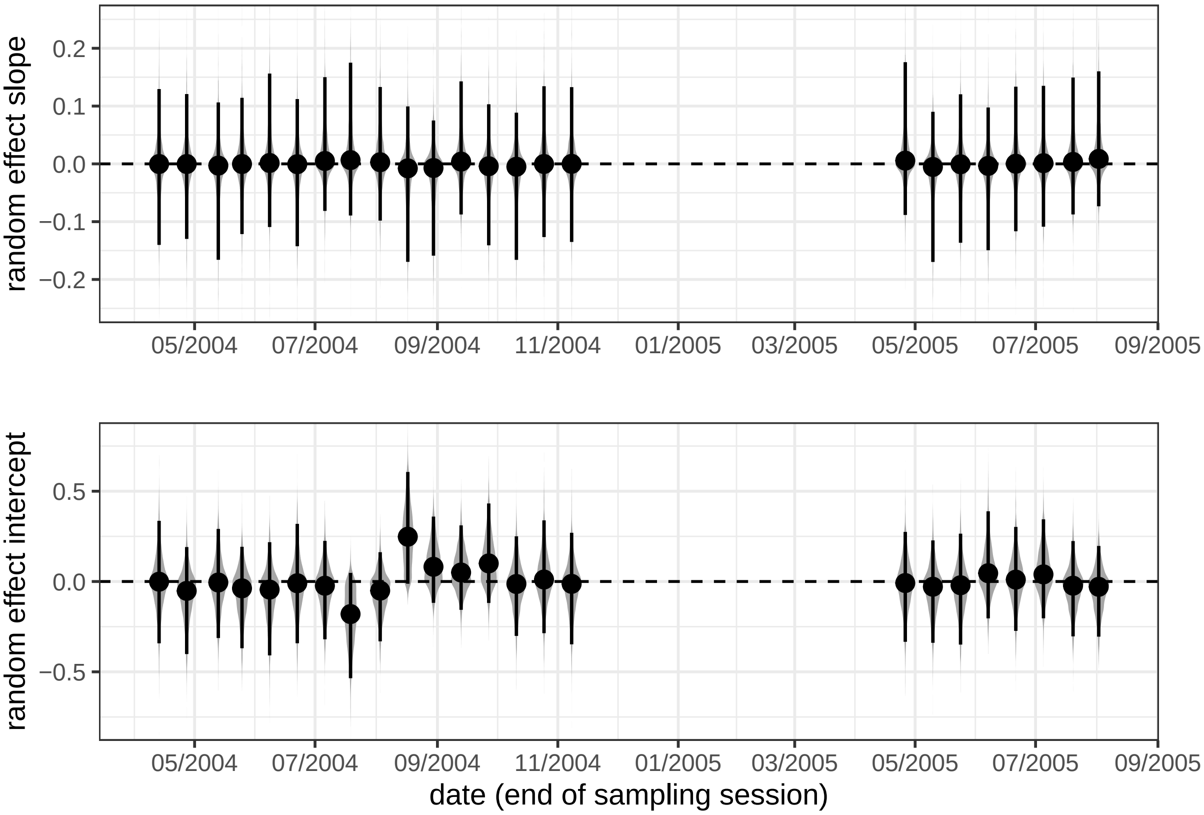


**Figure S4-1.** Posteriors for the session-specific random effect slopes (top) and intercepts (bottom), i.e. $\eta_j$ and $\gamma_j$ in the model description in **S1**, respectively. The model used is the beta-binomial model with Imperviousness Density (100 m) as the urbanisation metric. Black dots are posterior means, segments 95% Highest Posterior Density Intervals.

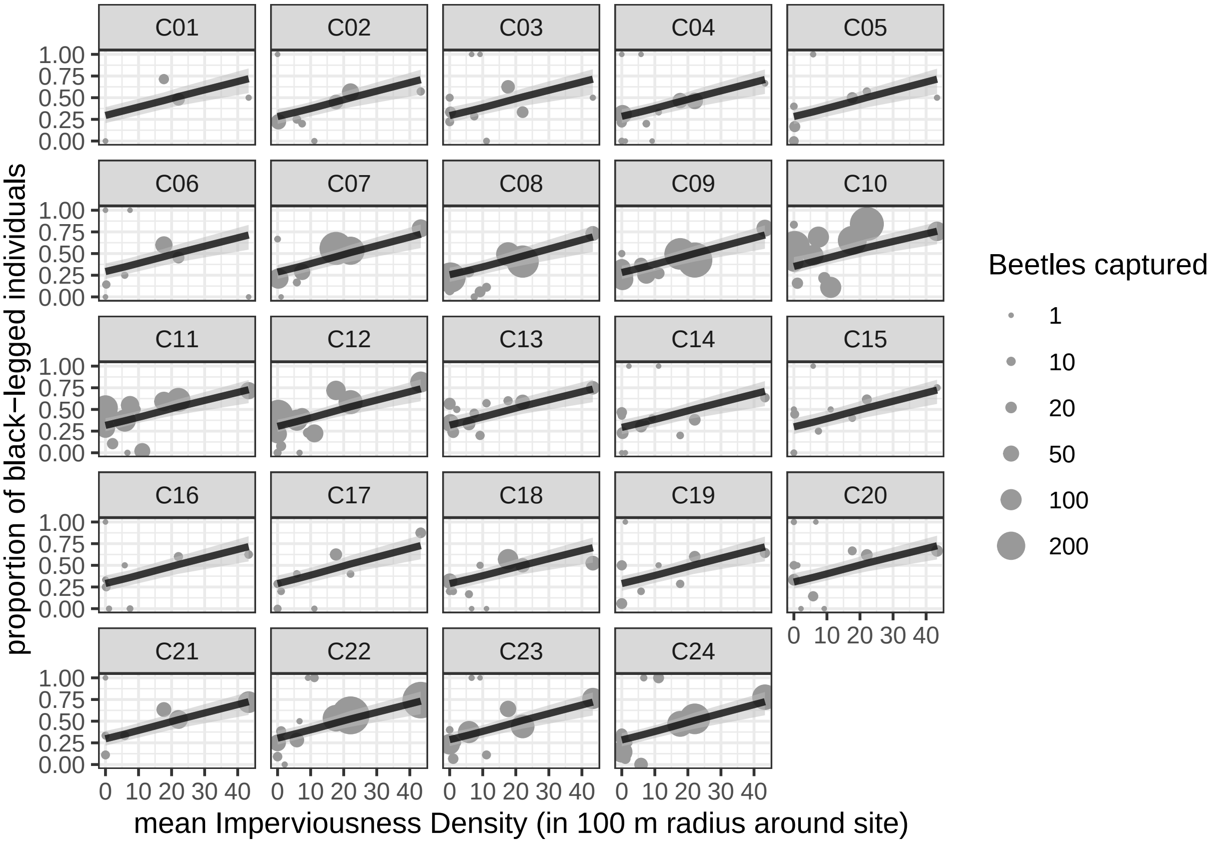


**Figure S4-2.** Proportion of *Pterostichus madidus* beetles with black legs as a function of urbanisation (Imperviousness Density in 100 m radius), displayed session by session. Grey dots: observed proportions, black line and grey bands: mean predicted relationship *at the sampling session level* (i.e. including random effects in the predictions) along with 95% credible bands (using Highest Posterior Density intervals).

### S5 - Population size variability along the urbanisation gradient

Evolutionary clines may in some cases be caused by non-adaptive forces; this includes genetic drift, in particular if a gradient in population sizes is correlated with the environmental gradient of interest. For instance in our case, if urbanisation leads to reduced population sizes (Santangelo et al. 2018). For a first check on whether our results may be explainable by drift, we looked at whether there was an effect of urbanisation on population sizes of *P. madidus* in our data.

Our models are here essentially structured the same way as the main proportion models (see main text and **S1**), with a few nuances summed up here:

- the response is now $N_{i,j}$ the total number of *P. madidus* caught by woodland $i$ × session $j$ combination;
- importantly, samples $ij$ with 0 beetles found are **included** here and not excluded;
- since data are now counts and not proportions, we use negative binomial models (with a log link) rather than (beta-)binomial ones;
- we include an offset or rate term to account for the fact sampling sessions are not the same length;
- the prior for the intercept $\beta_0$ is changed back to the "standard" $\text{Normal}(0, 1)$ from $\text{Normal}(0, 1.5)$, as the rationale for the latter is based on proportion data (McElreath 2020). All other priors are left as is.

Keeping the model structure of the main models (with random effects of sessions) allows us to account correctly for the known seasonality of beetle abundances (visible **Fig. S4-2**), rather than ignoring it by complete pooling, which may lead to potential issues.

We find that independently of the metric used, there is no clear evidence that urbanisation has an effect on population sizes (**Fig. S5-1**). We note that if there were to be an effect, the data suggests it would be towards *increased*, not reduced, population sizes in cities.

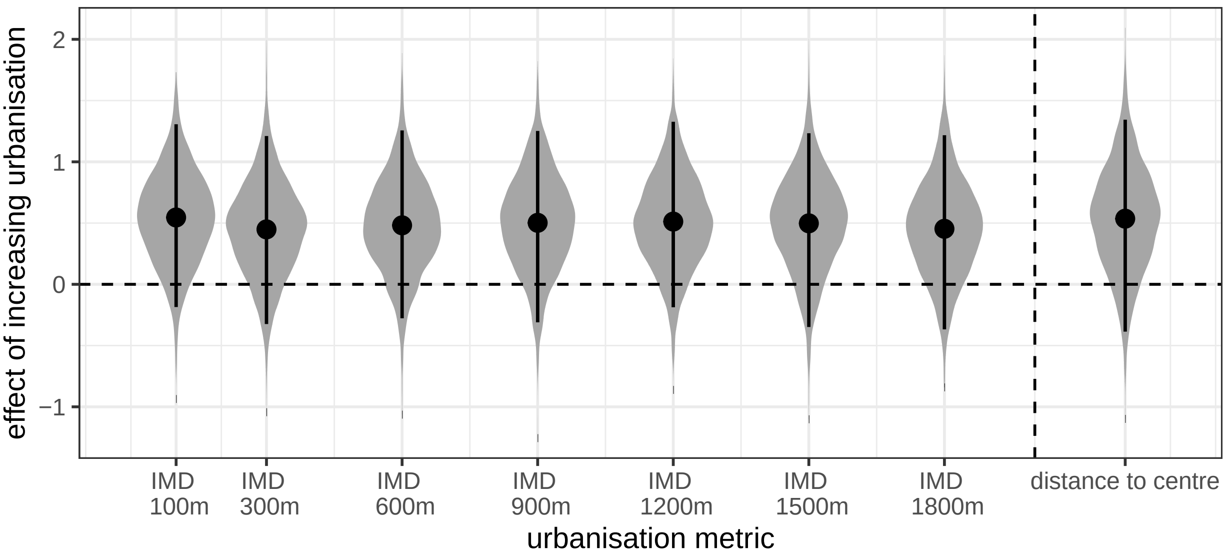


**Figure S5-1.** Posterior estimates for the standardised fixed effect of urbanisation on the number of *P. madidus* caught per day of sampling, depending on the urbanisation metric used in the model. Note that the sign of the posterior values for distance to city centroid has been reversed to make the comparison with the other posteriors easier (as distances to city centroid decrease when urbanisation increases). Black dots are posterior means, segments 95% Highest Posterior Density Intervals.

### References

Bürkner P-C (2017) Brms: An R package for Bayesian multilevel models using Stan. Journal of Statistical Software 80:1–28. https://doi.org/10.18637/jss.v080.i01

Harrison XA (2015) A comparison of observation-level random effect and Beta-Binomial models for modelling overdispersion in Binomial data in ecology & evolution. PeerJ 3:e1114. https://doi.org/10.7717/peerj.1114

McElreath R (2020) Statistical rethinking: A Bayesian course with examples in R and Stan, 2nd edition. Chapman and Hall/CRC, Boca Raton, USA

Nakagawa S, Schielzeth H (2013) A general and simple method for obtaining $R^2$ from generalized linear mixed-effects models. Methods in Ecology and Evolution 4:133–142. https://doi.org/10.1111/j.2041-210x.2012.00261.x

Santangelo JS, Johnson MTJ, Ness RW (2018) Modern spandrels: The roles of genetic drift, gene flow and natural selection in the evolution of parallel clines. Proceedings of the Royal Society B: Biological Sciences 285:20180230. https://doi.org/10.1098/rspb.2018.0230

Vehtari A, Gelman A, Gabry J (2017) Practical Bayesian model evaluation using leave-one-out cross-validation and WAIC. Statistics and Computing 27:1413–1432. https://doi.org/10.1007/s11222-016-9696-4
